# Predictors of intestinal inflammation in asymptomatic first-degree relatives of patients with Crohn’s disease

**DOI:** 10.1101/173492

**Authors:** Kirstin M. Taylor, Ken B. Hanscombe, Raquel Iniesta, Matthew Traylor, Nicola S. Taylor, Nicholas Powell, Peter M. Irving, Simon H. Anderson, Natalie J. Prescott, Christopher G. Mathew, Cathryn M. Lewis, Jeremy D. Sanderson

## Abstract

**Objective:** Relatives of individuals with Crohn’s disease (CD) carry an increased number of CD-associated genetic variants and are at increased risk of developing the disease. Multiple environmental and genetic factors contribute to this increased risk. We aimed to estimate the utility of genotype, smoking, family history, and a panel of biomarkers to predict risk in asymptomatic first-degree relatives (FDRs) of CD patients.

**Design:** We calculated a combined genotype (72 CD-associated genetic markers) and smoking relative risk score in 454 FDRs, and performed capsule endoscopy and collected 22 biomarkers in individuals from the highest and lowest risk quartiles. We then predicted small intestinal inflammation using genetic risk score, smoking status, number of relatives with CD, capsule transit time, and the panel of biomarkers in 124 individuals with complete data. Our principal analysis was to calculate the predictive utility from two machine learning classifiers: an elastic net and a random forest.

**Results:** Both classifiers successfully predicted FDRs with intestinal inflammation: elastic net (AUC=0.80, 95% CI: 0.62-0.98), random forest (AUC=0.87, 95% CI: 0.75-1.00). The elastic net selected a 3-predictor solution: CD family history (OR=1.31), genetic risk score (OR=1.14), and faecal calprotectin (OR=1.04). The same 3 variables were among the top 5 most important predictors as ranked by the random forest.

**Conclusion:** A readily collectable panel of genetic risk variants, added to family history and faecal calprotectin, predicts those at greatest risk for developing CD with a good degree of accuracy.

## Introduction

Inflammatory bowel disease (IBD), comprising Crohn’s disease (CD) and ulcerative colitis (UC), is a chronic inflammatory condition of the gastrointestinal tract associated with significant morbidity. Family members of individuals with CD are at increased risk for developing the disease. Estimates of the sibling relative risk (the ratio of disease risk in siblings compared with the rate in the general population) range from 15 to 42^1^. A retrospective study of the entire Danish population over a period of more than 40 years found that about 12% of new incidents of CD occurred in families already affected by the disease^2^. Disease onset precedes the development of symptoms, which often precede diagnosis by months to years, at which stage many patients have developed complications of the disease (including malnutrition, osteoporosis, and strictures or fistulas requiring surgery)^3^. There is emerging evidence to support the use of immunosuppressive treatment early in the course of CD to reduce the risk of such complications^4^. With an increasing incidence and prevalence of IBD worldwide^5^, a clinical tool that facilitates early detection of those at greatest risk opens up the possibility for early intervention to alter or halt aberrant immune and inflammatory responses before the development of overt disease, and perhaps, ultimately, disease prevention.

Although CD pathogenesis remains incompletely understood, genetic risk for the disease is well-established. Heritability, the proportion of population trait variance explained by genetic factors, estimated from identical and non-identical twin concordance rates, is 0.75^6^. Genetic variance explained by CD-associated loci discovered through genome-wide association studies (GWAS), so-called SNP heritability, is 0.37, indicating that approximately half the total CD heritability is explained by known SNPs^6^. To date, there are 240 single nucleotide polymorphisms (SNPs) robustly associated with the IBD^7^. A recent study of first-degree relatives (FDRs) of patients with IBD showed that they are enriched for IBD-associated risk loci^8^.

Familial clustering of disease is likely the result of shared environmental factors, as well as genetic risk factors. The observed increasing incidence of the disease among populations with historically lower rates, and those migrating from regions with low incidence rates to regions with higher rates, is consistent with a substantial environmental risk component to CD^5^. Many lifestyle-related factors have been implicated, including stress, sedentary lifestyle, western diet, poor sleep, and tobacco use^9 10^. Smoking is the best-studied environmental risk factor; a meta-analysis estimated a two-fold risk increase for CD in current smokers^11^.

Asymptomatic FDRs may display phenotypic features in common with CD patients including altered intestinal permeability, positive serological antimicrobial markers, disordered innate and acquired immunity, faecal dysbiosis, and elevated faecal calprotectin (FC)^12^. Overt small intestinal (SI) inflammation has been described in FDRs who have undergone ileocolonoscopy^13^, intestinal ultrasound^14^ and video capsule endoscopy (VCE)^15^.

Evidence of increased risk factors for CD in FDRs raises the possibility of predicting those at risk of developing the disease^12^. We hypothesized that the observed increase in the number of CD-associated risk loci in FDRs relative to healthy controls^8^, elevated levels of FC^12^, and smoking status, taken together could provide sufficient information to detect at-risk individuals before the development of overt symptoms. In this study we aimed to assess the clinical utility of a disease risk model to predict the presence of SI inflammation as detected by VCE. We applied two machine learning methods to genetic risk (72 genetic markers), smoking status, number of relatives with CD, 22 biomarkers, and assessed the predictive ability of the derived models.

## Material and Methods

### Participants

We recruited 480 healthy FDRs (full siblings, offspring or parents) between the age of 18 and 55, through patients with CD attending the IBD service at Guy’s & St Thomas’ NHS Foundation Trust (GSTT) and members of Crohn’s & Colitis UK (a charity supporting patients with IBD). CD diagnosis in probands was confirmed by their gastroenterologist or general practitioner. Interested FDRs provided information regarding family history of IBD, medical history (including gastrointestinal symptoms), smoking status, medication use, allergies, primary care provider and ethnicity. In keeping with the population of the meta-analysis of CD GWAS used for calculating the genotype relative risk^16^, only FDRs of European ancestry were included. The inclusion criteria were: ability to give written informed consent, age 18-55 years, confirmed history of Crohn’s disease in the proband, and absence of gastrointestinal symptoms in the FDR. The exclusion criteria were: a previous diagnosis of IBD, a previous diagnosis of irritable bowel syndrome, or presence of a major co-morbidity. Additional exclusion criteria for capsule endoscopy were: pregnancy, major bowel surgery, or the use of non-steroidal anti-inflammatory drugs in the 4 weeks prior to capsule endoscopy (low-dose aspirin excluded).

### DNA collection and genotyping

480 FDRs met the inclusion criteria and were sent a saliva sampling kit (Oragene^TM^ DNA OG-500) and 455 returned saliva samples. DNA extraction and genotyping protocols are included in the supplementary materials. DNA samples were genotyped on the Immunochip, which is a custom Illumina Infinium array containing 196,524 SNPs and small insertion/deletions selected mainly from GWAS analysis of 12 immune-mediated diseases^17^. It included all SNPs from CD-associated loci at the time of design. The three major *NOD2* variants were analysed separately (rs2066844, rs2066845 and rs2066847), instead of the *NOD2* tagging SNP, rs2076756. Owing to the enhanced risk of homozygotes and compound heterozygotes for *NOD2* variants, the *NOD2* variants were combined, rather than treated as independent loci. Genotypes for a total of 72 SNPs from known CD risk loci were extracted for genetic risk profiling.

The relevant SNP data were extracted and underwent quality control (QC) in PLINK^18^. SNPs with missingness >3% and Hardy-Weinberg equilibrium outliers (p ≤ 1 x 10^−4^) were excluded. X chromosome heterozygosity rates were used to determine sex empirically. Population structure was assessed by principal components analysis in PLINK as previously described^19^, derived from CEU HapMap 3 individuals (Northern and Western European Ancestry). No outliers were identified.

Following QC, 11 saliva samples were repeated due to a low DNA concentration or quality, one of which repeatedly failed QC and was excluded. 454 FDRs were successfully genotyped (61% female; median age 34, range 18-55; 40% were siblings, 46% offspring, 14% parents of probands; 44% were current or ex-smokers).

### Calculation of high and low risk groups

We generated a combined smoking and genotype risk score in 454 FDRs of individuals with CD using the R package REGENT^22^. Summary results were obtained for 72 genetic variants from GWAS meta-analysis in CD^16^. Smoking risk was calculated using ORs from a large case-control study^23^. 147 FDRs in the highest and lowest risk quartiles of the risk score were invited to undergo VCE, and to provide stool and blood samples for biomarker analysis.

Subsequent statistical analyses were performed on the subset of 124 participants for whom we had complete data on all measures (age in years (mean=37.42, sd=10.46), sex (male=42, female=82), and genetic risk score and smoking were considered in the modelling as separate variables.

Smoking status was recorded as "Current", "Ex", or "Never" smoked (29, 33, 62 respectively among 124 individuals with complete data). We used "Never" smoked as the reference level. The number of family members with CD (CD family history) was coded as "single" or "multiple" (111, 13 respectively among 124 individuals with complete data).

### Video capsule endoscopy (VCE)

Capsule endoscopies were performed in the Endoscopy Unit at GSTT following written informed consent. The MiroCam^tm^ (Intromedic, Seoul, Korea) VCE system was used for all capsule endoscopies (see supplementary materials for full description and protocol). Two validated scoring systems were used to quantify the degree of inflammation within the small intestine: the Lewis Score^20^ and the Capsule Endoscopy Crohn’s Disease Activity Index (CECDAI)^21^. SI transit time was defined as the passage from the first duodenal to the first caecal image, in minutes.

### Biomarkers

Full biomarker description and protocol are included in the supplementary materials.Stool was analysed for faecal calprotectin and serum for high-sensitivity C-reactive protein (hs-CRP), anti-saccharomyces cerevisiae antibodies (ASCA), and cytokines and growth factors. Table 1 shows the descriptive statistics for the complete list of 22 biomarkers measured. We dropped from the statistical analyses, 6/22 biomarkers with near zero variance among the 124 individuals with complete data.

**Table 1.**
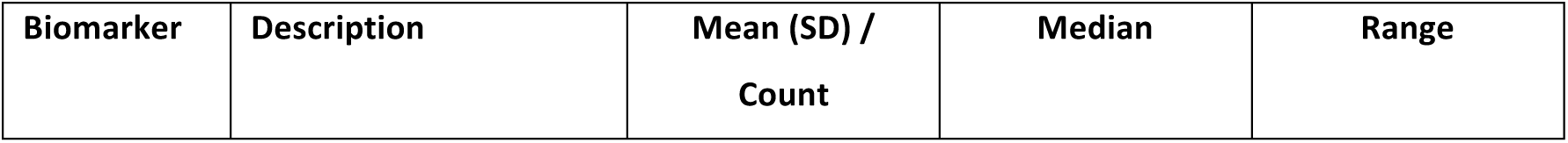
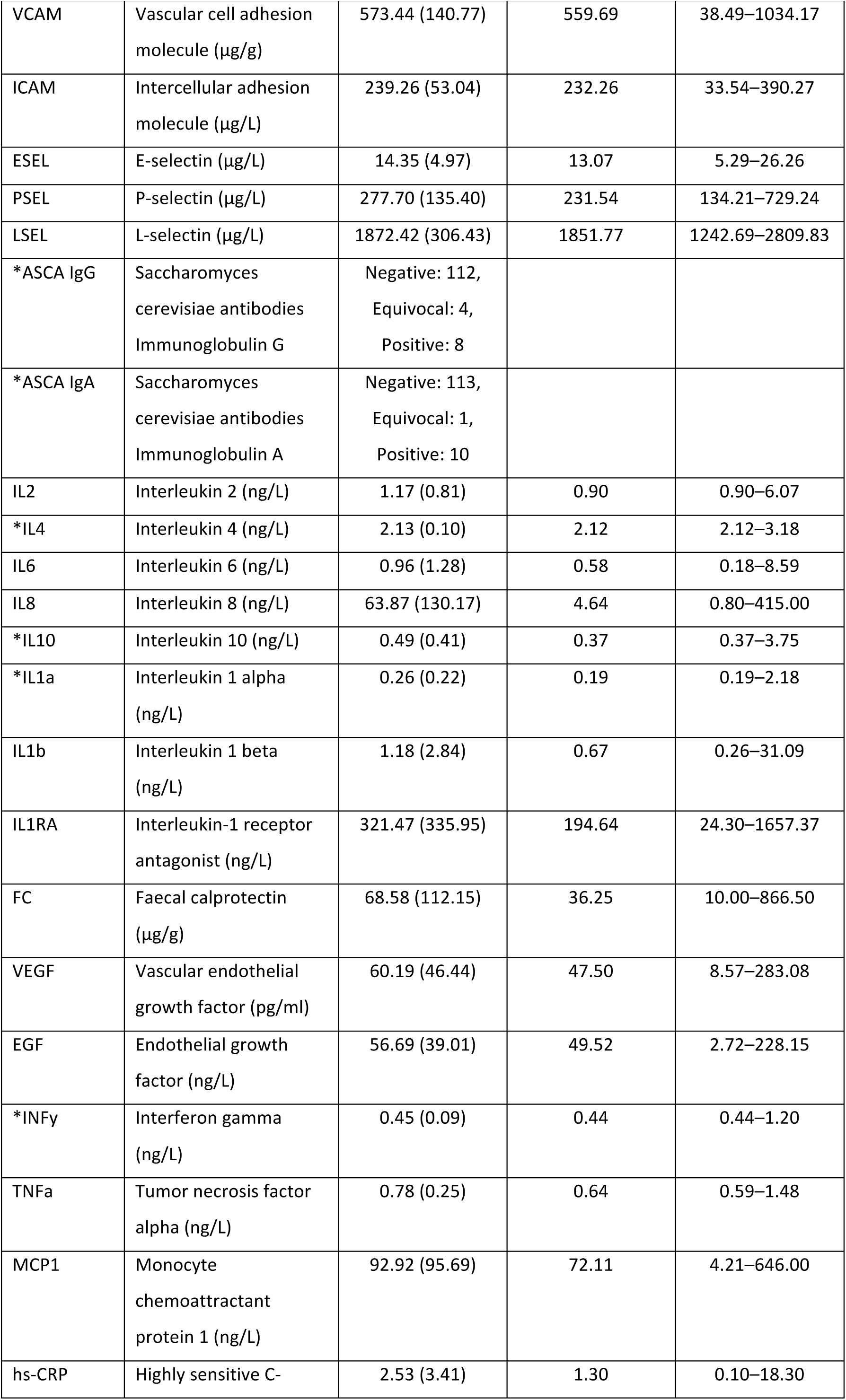
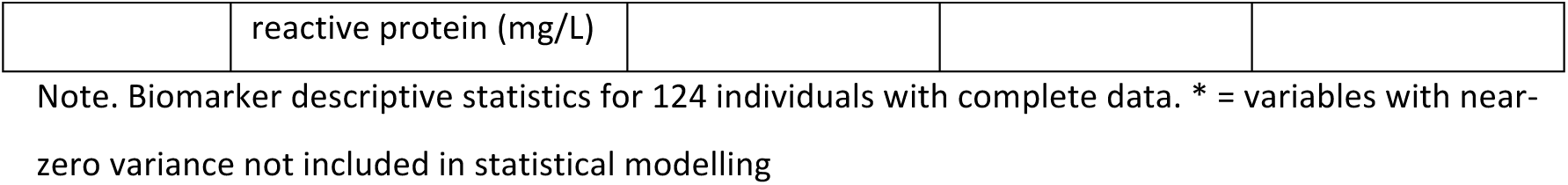
Biomarker descriptive statistics

| Biomarker | Description | Mean (SD) /<br>Count | Median | Range |
| --- | --- | --- | --- | --- |
| VCAM | Vascular cell adhesion molecule (µg/g) | 573.44 (140.77) | 559.69 | 38.49–1034.17 |
| ICAM | Intercellular adhesion molecule (µg/L) | 239.26 (53.04) | 232.26 | 33.54–390.27 |
| ESEL | E-selectin (µg/L) | 14.35 (4.97) | 13.07 | 5.29–26.26 |
| PSEL | P-selectin (µg/L) | 277.70 (135.40) | 231.54 | 134.21–729.24 |
| LSEL | L-selectin (µg/L) | 1872.42 (306.43) | 1851.77 | 1242.69–2809.83 |
| *ASCA IgG | Saccharomyces cerevisiae antibodies Immunoglobulin G | Negative: 112,<br>Equivocal: 4,<br>Positive: 8 |  |  |
| *ASCA IgA | Saccharomyces cerevisiae antibodies Immunoglobulin A | Negative: 113,<br>Equivocal: 1,<br>Positive: 10 |  |  |
| IL2 | Interleukin 2 (ng/L) | 1.17 (0.81) | 0.90 | 0.90–6.07 |
| *IL4 | Interleukin 4 (ng/L) | 2.13 (0.10) | 2.12 | 2.12–3.18 |
| IL6 | Interleukin 6 (ng/L) | 0.96 (1.28) | 0.58 | 0.18–8.59 |
| IL8 | Interleukin 8 (ng/L) | 63.87 (130.17) | 4.64 | 0.80–415.00 |
| *IL10 | Interleukin 10 (ng/L) | 0.49 (0.41) | 0.37 | 0.37–3.75 |
| *IL1a | Interleukin 1 alpha (ng/L) | 0.26 (0.22) | 0.19 | 0.19–2.18 |
| IL1b | Interleukin 1 beta (ng/L) | 1.18 (2.84) | 0.67 | 0.26–31.09 |
| IL1RA | Interleukin-1 receptor antagonist (ng/L) | 321.47 (335.95) | 194.64 | 24.30–1657.37 |
| FC | Faecal calprotectin (µg/g) | 68.58 (112.15) | 36.25 | 10.00–866.50 |
| VEGF | Vascular endothelial growth factor (pg/ml) | 60.19 (46.44) | 47.50 | 8.57–283.08 |
| EGF | Endothelial growth factor (ng/L) | 56.69 (39.01) | 49.52 | 2.72–228.15 |
| *INFy | Interferon gamma (ng/L) | 0.45 (0.09) | 0.44 | 0.44–1.20 |
| TNFa | Tumor necrosis factor alpha (ng/L) | 0.78 (0.25) | 0.64 | 0.59–1.48 |
| MCP1 | Monocyte chemoattractant protein 1 (ng/L) | 92.92 (95.69) | 72.11 | 4.21–646.00 |
| hs-CRP | Highly sensitive C- reactive protein (mg/L) | 2.53 (3.41) | 1.30 | 0.10–18.30 |

Note. Biomarker descriptive statistics for 124 individuals with complete data. \* = variables with near-zero variance not included in statistical modelling

### Statistical analysis

All statistical analyses were performed in the R programming language and environment^24^. For the predictive modelling we used the R package caret^25^, and its dependencies glmnet^26^ and ranger^27^.

### Explanatory modelling

We first fitted multivariate logistic regression models with dichotomized Lewis score (<135 = Normal i.e. without intestinal inflammation, ≥135 = Abnormal, i.e. with intestinal inflammation) as response variable, and age, sex, genetic risk score, smoking status, CD family history, VCE capsule transit time and the biomarkers as explanatory variables. Applying generalised linear regression models to a dataset with a large number of predictor variables relative to the number of samples is not optimal for several reasons including over-fitting (explaining noise as well as signal in the sample), multi-collinearity (correlation and potential redundancy among predictors), and low interpretability^28^. We used stepwise model selection to address this overfitting, multicollinearity, and model complexity. At each step in the iterative selection procedure, a variable is considered for addition to (or subtraction from) the current set of explanatory variables based on a model comparison criterion. We compared two criteria: Akaike’s information criterion (AIC) and the Bayesian information criterion (BIC). However, this approach does not eliminate the problems of overfitting, multicollinearity, and model complexity, especially with a small sample and large number of measured variables. In addition, explanatory models can quantify the relationship between predictors and outcome in the particular sample collected (e.g. regression coefficients, and total variance explained), but they give no measure of the predictive performance of the putative predictors in unseen cases. Given that the ultimate aim of the study was to estimate the utility of the genetic, environmental and biomarker variables to identify high risk individuals, we went on to derive a series of predictive models. Machine learning is specifically designed to deal with the large number of variables relative to sample size problem, and includes techniques to address the low events-per-variable ratio (small number of cases).

### Predictive modelling

Our goal was to develop a predictive model for SI inflammation, achieving a balance between interpretability (by reducing the number of predictor variables) and predictive ability. Thus we used machine learning, which has techniques for variable subset selection and estimation of how well a given model will perform at predicting future data^29^. Machine learning finds structure in data and addresses over-fitting when there are a large number of predictors. More generally, it is a set of techniques that improve performance on a specified task (e.g. classifying absence/presence of inflammation) with experience (exposure to data). We divided the data into a *training* sample (2/3) for model building and a *test* sample (1/3) for evaluation of model predictive performance and compared two machine learning techniques:

#### Elastic net

The elastic net is an extension of the basic regression framework that allows selection of the most important subset of predictors. It mixes a ridge penalty (which shrinks the coefficients of correlated predictors towards each other) and a LASSO penalty (which selects one among a group of correlated predictors and shrinks the coefficients of the others to zero) to perform variable selection^28^. We used 20 repeats of 5-fold cross validation to estimate model parameters (penalty mixing factor (α) and penalty strength (λ)) that optimised the model’s prediction performance, i.e., its ability to correctly classify individuals with and without inflammation. We measured prediction performance with the area under that receiver-operator characteristic curve (AUC: a plot of the true positive rate against the false positive rate for different cut-offs of the model estimated probability of being Abnormal) and accuracy (the fraction of test sample predictions that are true). The absolute value of the *t*-statistic for each parameter in the model is used to judge relative variable importance.

#### Random forest

A random forest is a collection of classification or regression trees. A tree is a series of splitting rules. At each split in a particular tree, from a random subset of predictors a single predictor is chosen that produces branches with the best split of Normal and Abnormal samples. Each resulting branch is then split until it ends in a "leaf" containing only (or mainly) Normal or Abnormal training samples. A test sample is then pass through each tree, and is assigned the (majority) class of the leaf on which it lands. Every tree in the forest produces a prediction and these predictions are combined to give a single consensus prediction for the individual^28 29^. We used 20 repeats of 5-fold cross validation on the training data to select the optimum number of randomly selected predictors (*mtry*) that maximized the AUC. Overall variable importance was determined by permuting predictors one at a time and measuring the mean decrease in accuracy averaged over all trees.

These two predictive modelling techniques, as well as more general statistical modelling and machine learning considerations including pre-processing, test-train dataset split, cross validation, variable importance, and class imbalance are described in detail in the supplementary material.

## Results

### Capsule endoscopy findings

147 FDRs underwent VCE: in 144 the caecum was reached. There was one capsule retention, managed conservatively with prokinetic agents. The commonest abnormal finding was of small aphthous ulceration in the distal ileum, and no strictures were identified. Marked inflammation typical of CD (>150 aphthous ulcers throughout the small intestine) was found in only one FDR, in the high risk group. 94% of the lowest risk group had no SI inflammation (Lewis score <135) compared with 68% of the highest risk group (p=0.0001). In the highest risk group, 9% had moderate-severe inflammation (Lewis score ≥790) compared with none in the low risk group (p=0.016).

### Characteristics of the highest and lowest risk quartiles

In total, 124 of the 147 FDRs who underwent VCE had complete study data and are included in all analyses. Table 2 shows the distribution of proband relationship, sex, age, smoking status, CD family history, and VCE-determined SI inflammation by combined genotype and smoking relative risk. Among these variables, only smoking and Lewis score showed a significant difference by risk quartile. The Lewis and CECDAI capsule scores were highly correlated (Pearson’s *r*=0.89, 95% CI=0.85-0.92, p<0.01) and when each measure was dichotomized (Normal, Abnormal) only 3 samples were classified differently. For all modelling we used the dichotomized Lewis score as our outcome, as this has previously been shown to correlate well with FC where the CECDAI did not^30^.

**Table 2.**
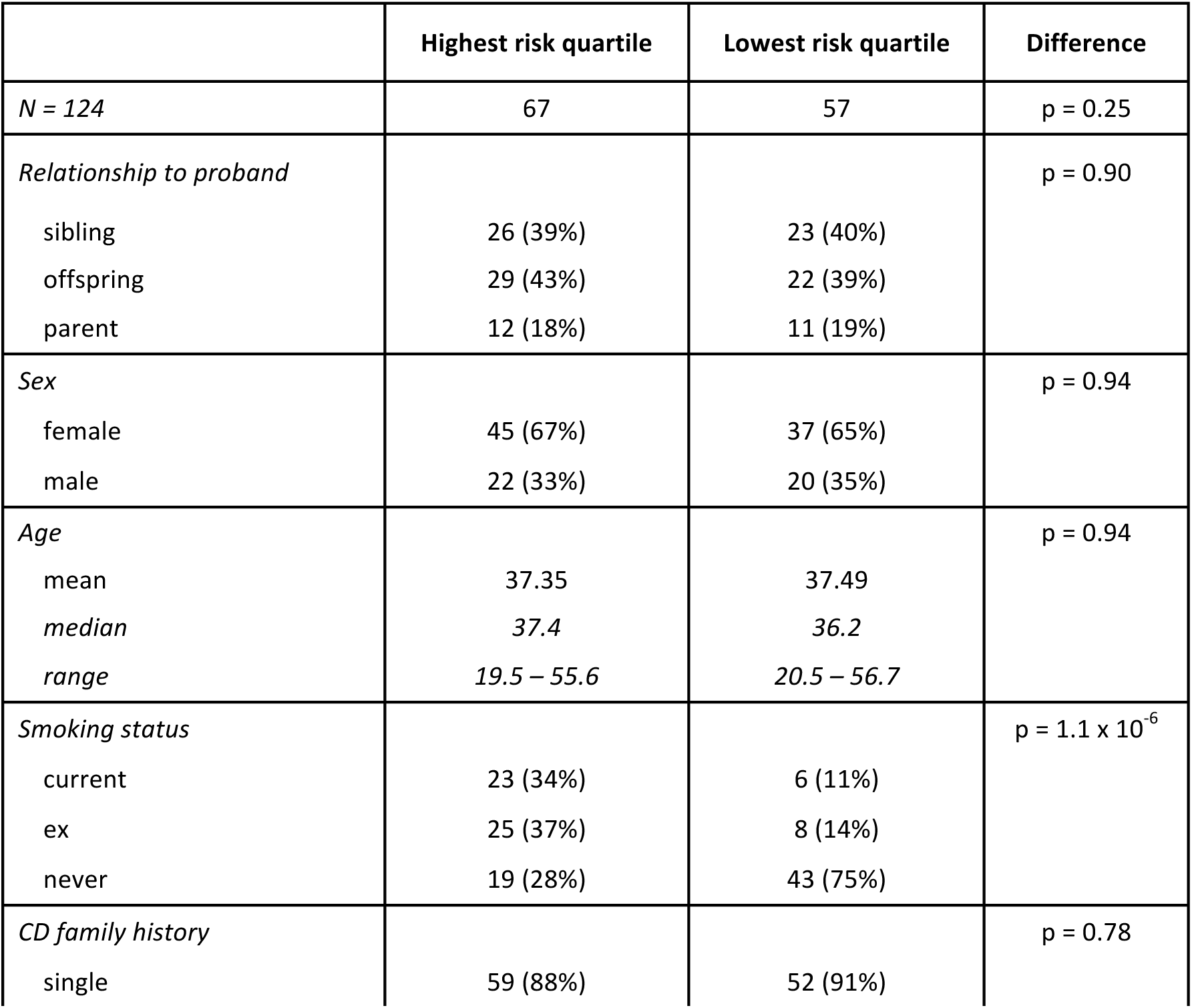
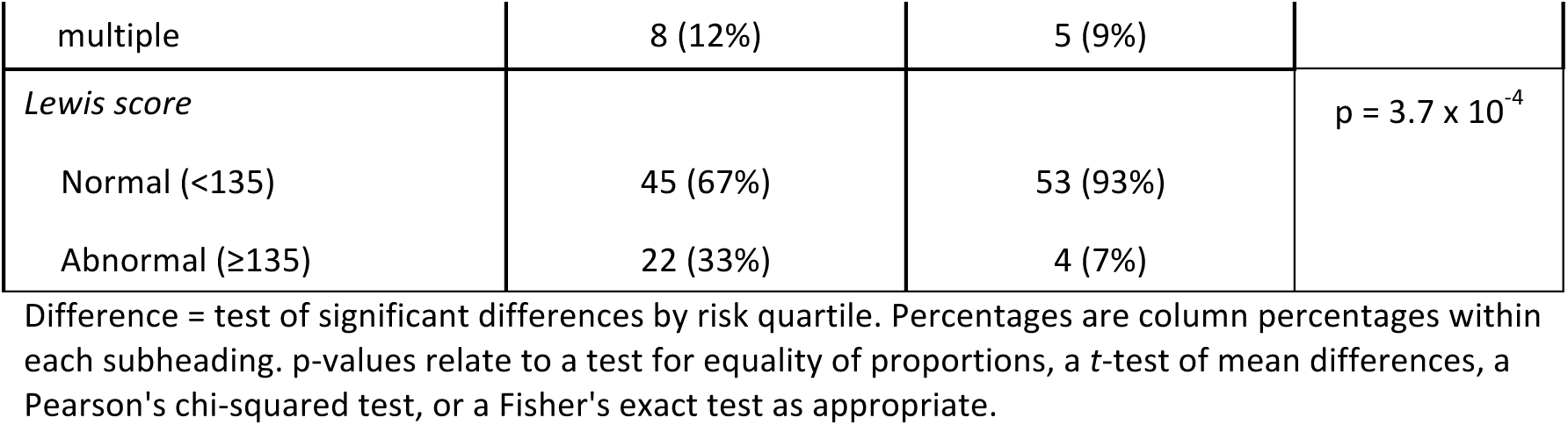
Distribution of demographics, relationship and Lewis score by risk quartile

### Explanatory modelling

A logistic regression model for SI inflammation (based on Lewis score) was built including all variables. Using stepwise selection and Akaike’s information criterion (AIC) we reduced the number of predictors to a best explanatory subset of 14 (R^2^=0.72). Stepwise selection using the Bayesian information criterion (BIC), which seeks a more parsimonious model, yielded a 3 variable model (R^2^=0.47) including FC, CD family history (binary-coded "single" or "multiple"), and genetic risk score (supplementary **Table S4**). Although stepwise selection achieves a reduction in the number of predictor variables, and can quantify the variance in the outcome explained by the predictors, these classic statistical approaches do not account for correlation among predictors (supplementary **Figure S1**) and provide no indication of the predictive ability of the derived model.

### Predictive modelling

Predictive models were built on a training sample of 83 FDRs (2/3 of the data), and predictive ability assessed on the test sample of 41 FDRs (remaining 1/3 of the data). The predictive performance of the random forest (AUC=0.87, 95% CI=0.75 – 1.00; Accuracy=0.73, 95% CI=0.57 – 0.86) was slightly better than the elastic net (AUC=0.80, 95%CI=0.62 – 0.98; Accuracy=0.68, 95% CI=0.52 – 0.82). This superior performance was the result of correctly classifying an additional 2 test samples (1 Normal, 1 Abnormal) among the 41 unseen test samples (Table 3). A full set of model performance metrics is included in supplementary **Table S5**.

**Table 3.**
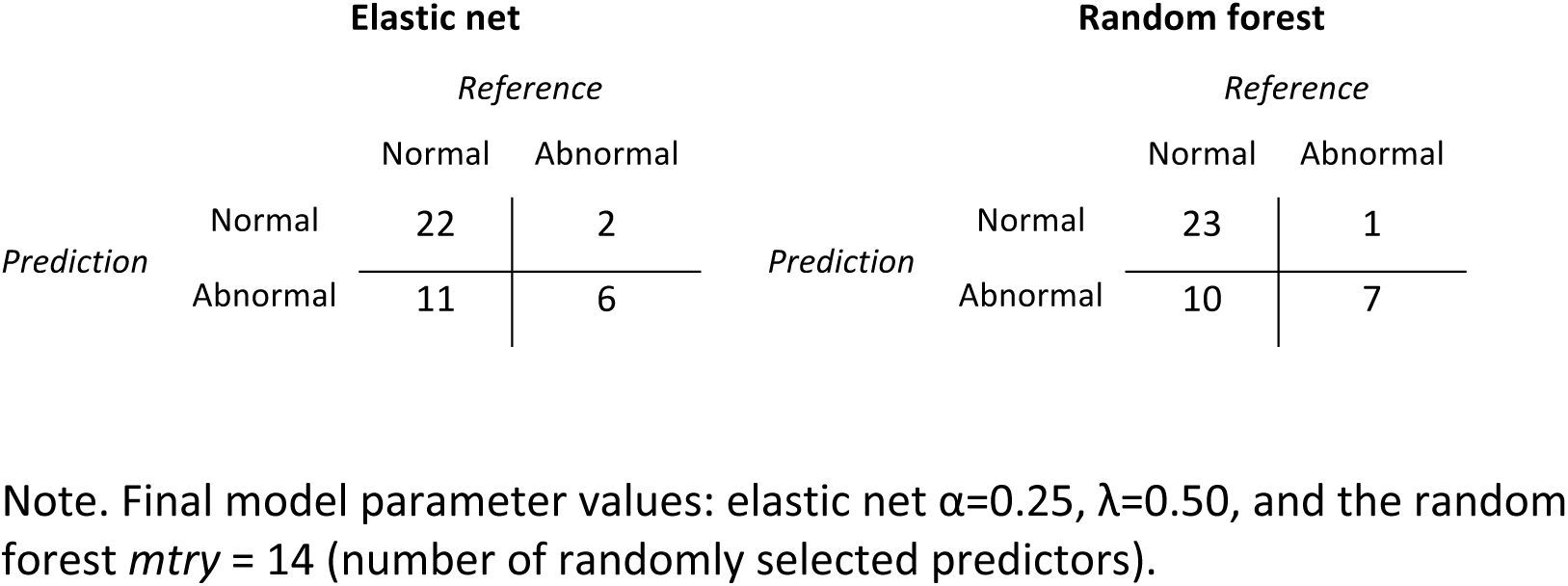
Confusion matrices: cross-tabulations of observed (reference) and predicted classification

| Elastic net |  |  |  | Random forest |  |  |  |
| --- | --- | --- | --- | --- | --- | --- | --- |
|  |  | Reference |  |  |  | Reference |  |
|  |  | Normal | Abnormal |  |  | Normal | Abnormal |
| Prediction | Normal | 22 | 2 | Prediction | Normal | 23 | 1 |
|  | Abnormal | 11 | 6 |  | Abnormal | 10 | 7 |

Note. Final model parameter values: elastic net $\alpha=0.25$ , $\lambda=0.50$ , and the random forest $mtry = 14$ (number of randomly selected predictors).

Figure 1 provides a visual summary of the predictive performance of our two classifiers. In a receiver-operator characteristic (ROC) curve, a good classifier moves away from the diagonal line (which indicates chance prediction) closer to the top left corner; in a precision-recall (PR) curve a good classifier bows closer to the top right hand corner. Taken together, the ROC and PR curves in Figure 1 suggest that the models performed similarly, and above chance.

**Figure 1.**
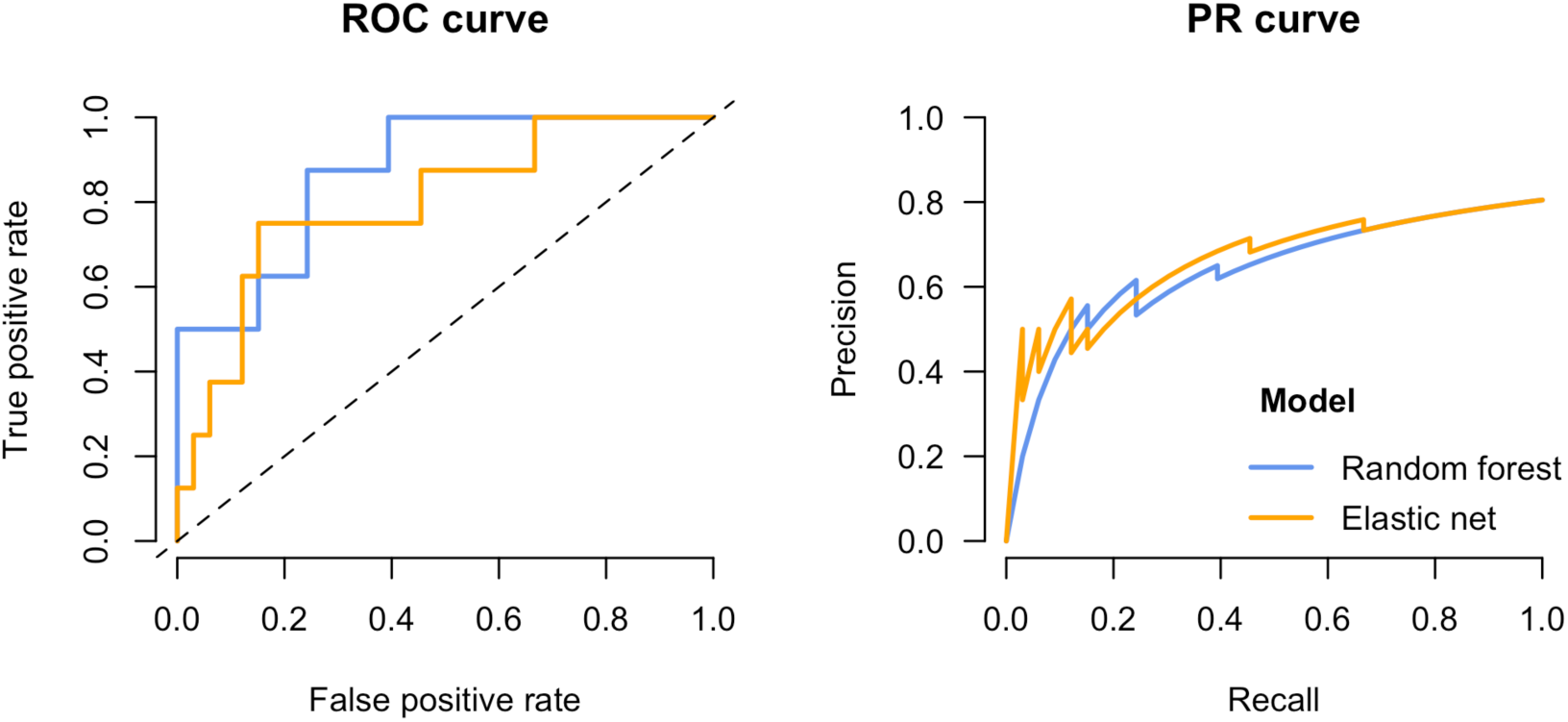
Classifier evaluation. Test sample performance was similar for the elastic net and random forest: elastic net sensitivity=0.75, specificity=0.67; random forest sensitivity=0.88, specificity=0.70. ROC curve: True positive rate (or Sensitivity) = TP / (TP+FN); True negative rate (or 1-Specificity) = 1 - (TN / (TN+FP)); PR curve: Precision (or Positive Predictive Value) = TP / (TP+FP) and incorporates the type I error rate; Recall = Sensitivity

### Predictor importance

Table 4 shows the predictors selected by the elastic net and their relative importance in both classifiers. The elastic net reduced the full set of predictors to CD family history (OR=1.31), genetic risk score (OR=1.14), and FC (OR=1.04), which are the same 3 predictors selected by stepwise BIC (supplementary **Table S4**). There is no acceptable way to assign p-values or confidence intervals to individual predictors in a cross-validated elastic net solution as standard approaches ignore "the complex selection procedure for defining the reduced model in the first place"^31^. Beyond the standard clinical predictors (family history and FC), genetic risk provided additional predictive utility.

**Table 4.**
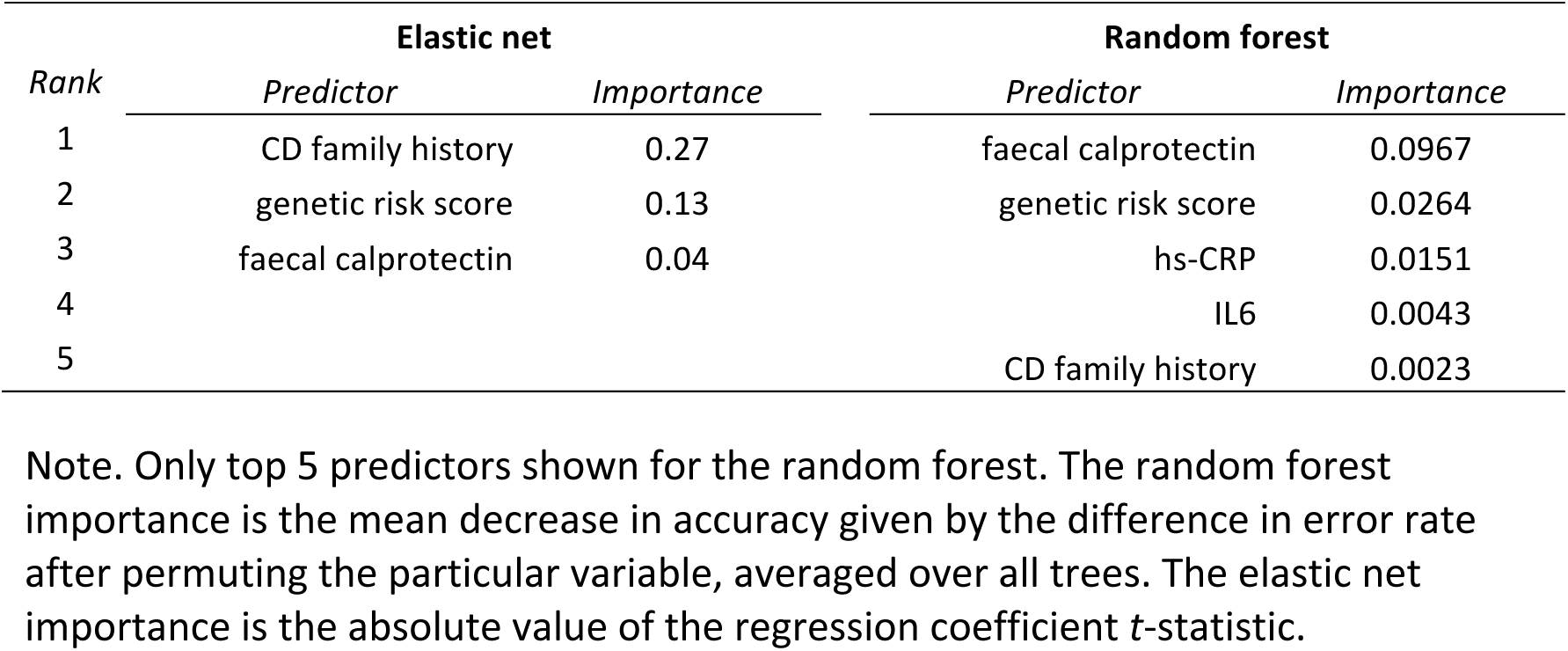
Relative predictor importance

### Genetic risk score improves predictive performance

Genetic risk score was one of three variables selected by the elastic net and ranked the second most important predictor by both classifiers. However, it is still relevant to determine how much additional predictive value genetic risk adds. A model built on all variables excluding genetic risk score produced a lower AUC (0.78, 95% CI: 0.55 – 1.00, p=0.0071) based on two variables: FC (OR=1.09) and CD family history (OR=1.63), compared to the full elastic net model AUC (0.80, 95% CI=0.62 – 0.98, p=0.0039 (Mann-Whitney *U* test derived^32^)).

## Conclusion

In our sample of 124 asymptomatic FDRs, we found that the combination of number of CD-affected relatives, genetic risk score, and level of FC made a good predictor of SI inflammation. The elastic net and random forest classifiers performed similarly, with significant above-chance prediction of SI inflammation.

The two models assume a different relationship between predictors and outcome: elastic net combines predictors linearly, whereas random forest makes multiple binary splits of the observations on one predictor at a time. In the elastic net, it is the combined set of selected predictors that captures the variance explained by correlated factors not selected. For example, the elastic net did not select smoking. This does not suggest smoking is any less important as a risk factor for the disease but is simply a product of the study design and the correlational pattern of the particular predictors included. The random forest, which by design does not exclude predictors, ranked smoking among the top 10 most important variables. Given the different design of the classifiers, there is no expectation that they should agree. However, that their results converge – CD family history, genetic risk score, and FC appeared among the top 5 predictors of the random forest – increases confidence in the utility of the particular set of predictors included.

CD family history was the strongest predictor in the elastic net model, and ranked 5^th^ most important predictor in the random forest. A retrospective study of the entire Danish population followed for a 44-year period found that the incidence rate for IBD was increased among individuals with two or more affected relatives^2^. By design, all our study participants had at least one relative with CD; we found that having a stronger family history (two or more relatives with CD) increased the risk for SI inflammation. It is worth noting that although studies often include the raw number of affected family members as a risk factor, it is a crude measure of familial risk. Ideally, total family size, structure, and age of affected individuals as well as raw count should be incorporated in a measure of "family history"^33^.

We found that 44 of 124 FDRs (35%) had abnormally elevated FC (≥50 μg/g, usually indicative of intestinal pathology^12^). Asymptomatic FDRs with increased FC have previously been observed^12^. Given that FC is part of the diagnostic work-up for CD^10^, and increased levels predict relapse^34^, our finding of the predictive utility of FC for SI inflammation suggest it is a promising biomarker for detecting those at greatest risk for CD. However, using a cut-off of ≥50 μg/g might include too many false positives - a cut-off of >250 μg/g may be a more appropriate level for a screening tool in asymptomatic individuals as this correlates with active CD^35^.

Genetic risk score was a significant predictor of SI inflammation. The predictive accuracy of our elastic net classifiers with or without genotype were both within the 0.7–0.8 range generally considered to be "acceptable discrimination"^36^. One way to evaluate the added value of genetic risk score is to consider what difference it might make in clinical practice. Our estimates showed a 2-percentage point increase in AUC on inclusion of genetic risk. Given that AUC is the probability that in a randomly selected case-control pair the case will be assigned a higher risk score, the model including genetic risk score accurately predicts an additional 2 cases in every 100 randomly selected pairs over the model without genetic risk. Updating the genetic risk panel as more risk loci are discovered is likely to increase the performance of the model further, and would be feasible now that low cost genome-wide SNP arrays are available which provide good coverage of known common IBD risk variants.

Mild SI inflammation found at VCE has been reported in 24% of asymptomatic FDRs, but the subsequent development of CD was not determined^37^. A study of 38 FDRs who underwent ileocolonoscopy found mild endoscopic and histological inflammation in 26% and CD in 13%. Those with mild inflammation were followed up with repeat ileocolonoscopy after a mean of 53 months without endoscopic or histological progression of inflammation^13^. In our study 26/124 (21%) of FDRs had abnormal Lewis scores, which is broadly consistent with these data. Long term follow-up of our FDR cohort is planned (out to 10 years), and may provide further information regarding risk of developing overt CD as opposed to asymptomatic SI inflammation. It is not yet certain whether these features are predictive of future development of CD, or are simply a spectrum of the "at-risk" individual who has not had the necessary environmental exposure(s) required to develop disease. Progression to CD in cases of isolated terminal ileitis found at ileocolonoscopy in the general population is not clear-cut, largely due to small study sample sizes, short duration of follow-up, heterogeneity of symptoms, and retrospective study design. Nevertheless, the rate appears to be low: 1% in a study with a mean follow-up of 29.9 months^38^,and 5% in another with a median follow-up of 97.5 months.^39^.

Although we used an unseen subset of samples (set aside from the full sample) to test the predictive performance of our model, this test sample was subject to the same design decisions as the training samples, as well as any study-specific idiosyncrasies. Ultimately our finding will require external validation, i.e., replication in a completely independent sample. We included 72 CD-associated genetic variants known at the time of clinical assessment^16^, and the single best-understood lifestyle factor, smoking. A replication could benefit from the inclusion of the up-to-date list of 240 IBD-associated SNPs^7^ (given that most IBD loci confer risk for both CD and UC^40^), and a more comprehensive picture of the environmental risk (i.e., in addition to smoking, medication, diet, stress, sleep, physical activity^9^). Finally, this study did not assess the gut microbiome, a likely predictor of risk for CD and potential target for intervention. The Genetic, Environmental and Microbiome (GEM) Project, currently recruiting 5000 CD FDRs internationally, will hopefully bring further insights into pre-clinical CD with the combination of all of these factors^41^.

In parallel with our increasing understanding of the specific genetic and environmental risk factors that combine to make an individual’s immune system hostile to commensal gut flora, a clinically useful tool for detecting those at greatest risk for CD before the presentation of overt symptoms would prioritise patient screening and follow up. Early detection opens up the possibility for early intervention, additional targets for drug development, and disease prevention. Our study suggests that a CD prediction tool can be built from a small set of biomarkers, known genetic risk variants, and family history of CD. Inclusion of risk factors from the most recent findings will improve prediction accuracy.

### Ethical approval & consent

Ethical approval was granted by Health Research Authority, UK (10/H0710/011) and all participants provided their written informed consent.

## References

1. Halme L, Paavola-Sakki P, Turunen U, et al. Family and twin studies in inflammatory bowel disease. World J Gastroenterol 2006;12(23):3668–72.

2. Moller FT, Andersen V, Wohlfahrt J, et al. Familial risk of inflammatory bowel disease: a population-based cohort study 1977-2011. Am J Gastroenterol 2015;110(4):564–71.

3. Thia KT, Sandborn WJ, Harmsen WS, et al. Risk factors associated with progression to intestinal complications of Crohn’s disease in a population-based cohort. Gastroenterology 2010;139(4):1147–55.

4. Khanna R, Bressler B, Levesque BG, et al. Early combined immunosuppression for the management of Crohn’s disease (REACT): a cluster randomised controlled trial. Lancet 2015;386(10006):1825–34.

5. Molodecky NA, Soon IS, Rabi DM, et al. Increasing incidence and prevalence of the inflammatory bowel diseases with time, based on systematic review. Gastroenterology 2012;142(1):46–54.e42; quiz e30.

6. Gordon H, Trier Moller F, Andersen V, et al. Heritability in inflammatory bowel disease: from the first twin study to genome-wide association studies. Inflamm Bowel Dis 2015;21(6):1428–34.

7. de Lange KM, Moutsianas L, Lee JC, et al. Genome-wide association study implicates immune activation of multiple integrin genes in inflammatory bowel disease. Nat Genet 2017;49(2):256–61.

8. Kevans D, Silverberg MS, Borowski K, et al. IBD Genetic Risk Profile in Healthy First-Degree Relatives of Crohn’s Disease Patients. J Crohns Colitis 2016;10(2):209–15.

9. Ananthakrishnan AN. Epidemiology and risk factors for IBD. Nat Rev Gastroenterol Hepatol 2015;12(4):205–17.

10. Baumgart DC, Sandborn WJ. Crohn’s disease. Lancet 2012;380(9853):1590–605.

11. Mahid SS, Minor KS, Soto RE, et al. Smoking and inflammatory bowel disease: a meta-analysis. Mayo Clin Proc 2006;81(11):1462–71.

12. Hedin CR, Stagg AJ, Whelan K, et al. Family studies in Crohn’s disease: new horizons in understanding disease pathogenesis, risk and prevention. Gut 2012;61(2):311–8.

13. Sorrentino D, Avellini C, Geraci M, et al. Tissue studies in screened first-degree relatives reveal a distinct Crohn’s disease phenotype. Inflamm Bowel Dis 2014;20(6):1049–56.

14. Biancone L, Calabrese E, Petruzziello C, et al. A family study of asymptomatic small bowel Crohn’s disease. Dig Liver Dis 2014;46(3):276–8.

15. Teshima CW, El-Kalla M, Turk SA, et al. Asymptomatic First Degree Relatives of Crohn’s Patients Display Endoscopic Small Intestinal Lesions Independent of Their Gut Permeability Status. Gastroenterology 2012;142(5):S–54.

16. Franke A, McGovern DP, Barrett JC, et al. Genome-wide meta-analysis increases to 71 the number of confirmed Crohn’s disease susceptibility loci. Nat Genet 2010;42(12):1118–25.

17. Cortes A, Brown MA. Promise and pitfalls of the Immunochip. Arthritis Res Ther 2011;13(1):101.

18. Purcell S, Neale B, Todd-Brown K, et al. PLINK: a tool set for whole-genome association and population-based linkage analyses. Am J Hum Genet 2007;81(3):559–75.

19. Anderson CA, Pettersson FH, Clarke GM, et al. Data quality control in genetic case-control association studies. Nat Protoc 2010;5(9):1564–73.

20. Gralnek IM, Defranchis R, Seidman E, et al. Development of a capsule endoscopy scoring index for small bowel mucosal inflammatory change. Aliment Pharmacol Ther 2008;27(2):146–54.

21. Gal E, Geller A, Fraser G, et al. Assessment and validation of the new capsule endoscopy Crohn’s disease activity index (CECDAI). Dig Dis Sci 2008;53(7):1933–7.

22. Crouch DJ, Goddard GH, Lewis CM. REGENT: a risk assessment and classification algorithm for genetic and environmental factors. Eur J Hum Genet 2013;21(1):109–11.

23. Brant SR, Wang MH, Rawsthorne P, et al. A population-based case-control study of CARD15 and other risk factors in Crohn’s disease and ulcerative colitis. Am J Gastroenterol 2007;102(2):313–23.

24. R Core Team. R: A language and environment for statistical computing. Secondary R Core Team. R: A language and environment for statistical computing 2013. http://www.R-project.org/.

25. caret: Classification and Regression Training. R package version 6.0-76 [program], 2017.

26. Friedman JH, Hastie T, Tibshirani R. Regularization Paths for Generalized Linear Models via Coordinate Descent. Journal of Statistical Software; Vol 1, Issue 1 (2010) 2010.

27. Wright MN, Ziegler A. ranger: A Fast Implementation of Random Forests for High Dimensional Data in C++ and R. Journal of Statistical Software; Vol 1, Issue 1 (2017) 2017.

28. James G, Witten D, Hastie T, et al. An introduction to statistical learning: with applications in R. New York: Springer, 2013.

29. Kuhn M, Johnson K. Applied Predictive Modeling. New York: Springer, 2013.

30. Koulaouzidis A, Douglas S, Plevris JN. Lewis score correlates more closely with fecal calprotectin than Capsule Endoscopy Crohn’s Disease Activity Index. Dig Dis Sci 2012;57(4):987–93.

31. Wu TT, Chen YF, Hastie T, et al. Genome-wide association analysis by lasso penalized logistic regression. Bioinformatics 2009;25(6):714–21.

32. Mason SJ, Graham NE. Areas beneath the relative operating characteristics (ROC) and relative operating levels (ROL) curves: Statistical significance and interpretation. Quarterly Journal of the Royal Meteorological Society 2002;128(584):2145–66.

33. Yasui Y, Newcomb PA, Trentham-Dietz A, et al. Familial relative risk estimates for use in epidemiologic analyses. Am J Epidemiol 2006;164(7):697–705.

34. Ikhtaire S, Shajib MS, Reinisch W, et al. Fecal calprotectin: its scope and utility in the management of inflammatory bowel disease. J Gastroenterol 2016;51(5):434–46.

35. D’Haens G, Ferrante M, Vermeire S, et al. Fecal calprotectin is a surrogate marker for endoscopic lesions in inflammatory bowel disease. Inflamm Bowel Dis 2012;18(12):2218–24.

36. Hosmer DW, Lemeshow S. Applied logistic regression. 2nd ed. New York ; Chichester: Wiley, 2000.

37. Teshima CW, Goodman KJ, El-Kalla M, et al. Increased Intestinal Permeability in Relatives of Patients With Crohn’s Disease Is Not Associated With Small Bowel Ulcerations. Clin Gastroenterol Hepatol 2017.

38. Chang HS, Lee D, Kim JC, et al. Isolated terminal ileal ulcerations in asymptomatic individuals: natural course and clinical significance. Gastrointest Endosc 2010;72(6):1226–32.

39. Tse CS, Deepak P, Smyrk T, et al. Sa1927 Natural History of Isolated Terminal Ileitis in Patients Without an Existing Diagnosis of Inflammatory Bowel Disease. Gastroenterology 2016;150(4):S406.

40. Cleynen I, Boucher G, Jostins L, et al. Inherited determinants of Crohn’s disease and ulcerative colitis phenotypes: a genetic association study. Lancet 2016;387(10014):156–67.

41. GEM Project. Crohn’s and Colitis Canada Inflammatory Bowel Disease. Secondary Crohn’s and Colitis Canada Inflammatory Bowel Disease. http://www.gemproject.ca/.

